# Gymnasts’ ability to modulate bioelectric sensorimotor rhythms during kinesthetic motor imagery of sports non-specific movements

**DOI:** 10.1101/2021.05.17.444451

**Authors:** Hirotaka Sugino, Junichi Ushiyama

**Author notes:** **Corresponding Author**: Junichi Ushiyama, Ph.D., Faculty of Environment and Information Studies, Keio University, 5322 Endo, Fujisawa, Kanagawa 252-0882, Japan.

## Abstract

Previous psychological studies using questionnaires have consistently reported that athletes have superior motor imagery ability, both for sports-specific and sports non-specific movements. However, regarding motor imagery of sports non-specific movements, no physiological studies have demonstrated differences in neural activity between athletes and non-athletes. The purpose of the present study was to examine differences in bioelectric sensorimotor rhythms during kinesthetic motor imagery (KMI) of sports non-specific movements between gymnasts and non-gymnasts. We selected gymnasts as an example population because they are likely to have particularly superior motor imagery ability due to frequent usage of motor imagery including KMI as part of daily practice. Healthy young participants (16 gymnasts and 16 non-gymnasts) performed repeated motor execution and KMI of sports non-specific movements (wrist dorsiflexion and shoulder abduction of the dominant hand). Scalp electroencephalogram (EEG) was recorded over the contralateral sensorimotor cortex. During motor execution and KMI, sensorimotor EEG power is known to decrease in the α- (8–15 Hz) and β-bands (16–35 Hz), referred to as event-related desynchronization (ERD). We calculated the maximal peak of ERD both in the α- (αERDmax) and β-bands (βERDmax) as a measure of changes in corticospinal excitability. αERDmax was significantly greater in gymnasts, who subjectively evaluated their KMI as being more vivid, for both KMI tasks. On the other hand, βERDmax was greater in gymnasts only for shoulder abduction KMI. These findings suggest gymnasts’ signature of flexibly modulating sensorimotor rhythm with no movements, which may be the basis of their superior ability of KMI for sports non-specific movements.

**New & Noteworthy:** Kinesthetic motor imagery of sports non-specific movements was compared between gymnasts and non-gymnasts (i.e., healthy controls) from both physiological and psychological approaches. The EEG sensorimotor rhythms during kinesthetic motor imagery were more desynchronized in gymnasts who subjectively imaged their own movements as being more vivid. The work reveals novel ability in gymnasts to flexibly control their sensorimotor rhythms with no actual movements, which would be the basis of their superior ability of motor imagery.

## Introduction

Motor imagery is regularly used by athletes to improve performance (1,2). Previous studies have shown that performing motor imagery in training improves performance in various tasks, including sequence learning (3,4), jump height (5) and free-throw shooting (6). To explain these performance gains, several neuroscience studies have provided evidence that motor imagery activates some neural substrates in common with actual movement, including the primary motor cortex, supplementary motor area, and inferior parietal lobe (7–10), as well as inducing neural plasticity in these areas (11–13).

In the field of sports psychology, several cross-sectional questionnaire studies have consistently reported that athletes have superior motor imagery ability compared with non-athletes (14–16). Furthermore, previous studies have indicated that athletes can perform motor imagery more vividly than non-athletes, not only for specialized movements in their own sports but also for sports non-specific movements such as raising the arm and jumping (15,17). The findings of these studies suggest that the neural activity underlying motor imagery ability may differ between athletes and non-athletes, not only for sports-specific imagery but also for motor imagery of sports non-specific movements.

However, in the field of applied physiology, to the best of our knowledge, no studies have demonstrated differences in neural activity during motor imagery of sports non-specific movements between athletes and non-athletes, although some studies using electroencephalogram (EEG) (17), magnetoencephalography(MEG) (18), functional magnetic resonance imaging (fMRI) (19) and transcranial magnetic stimulation (TMS) (20) have reported signature neural activity patterns during sports-specific motor imagery in athletes. For instance, when tennis players imagined movements specifically related to tennis, their corticospinal excitability became higher than that of non-athletes, whereas such a difference between athletes and non-athletes was not observed when they imagined other movements including non-tennis specific movements(20). Thus, there is currently a gap in findings between psychological and physiological studies regarding differences in motor imagery ability, particularly for sports non-specific movements between athletes and non-athletes.

To clarify this issue, the present study investigated differences in motor imagery ability of sports non-specific movements between gymnasts and healthy adults (i.e., non-gymnasts) from both psychological and physiological points of view. As a psychological indicator, we evaluated the subjective vividness of motor imagery using The Kinesthetic and Visual Imagery Questionnaire (KVIQ-20) (21). As a physiological indicator, we evaluated sensorimotor rhythms using EEG. Event-related desynchronization (ERD) is a measure of decreases in power of the EEG sensorimotor rhythm within the α- and β-bands from the resting-state to the motor execution or kinesthetic motor imagery (KMI) state (22,23), which is known to reflect increased neuronal excitability in the corticospinal system (24). We chose gymnasts as a population of athletes for the following reasons: 1) gymnasts perform motor imagery including KMI frequently as a part of their daily practices because of the high risk of serious injury in their performance; 2) gymnasts were assumed to have higher motor imagery abilities than athletes engaged in other sports, because it has been reported that motor imagery is more vivid in athletes engaged in individual and/or non-contact sports compared with athletes engaged in team and/or contact sports (25); 3) to the best of our knowledge, no previous studies have measured neural activity during motor imagery in gymnasts.

## Methods

### Ethical Approval

This study was conducted in accordance with the Declaration of Helsinki. All experimental protocols and procedures were approved by the Research Ethics Committee in Shonan Fujisawa Campus, Keio University (Approval Number 167). The examiners provided a detailed explanation of the purpose, experimental procedures, potential benefits, and risks involved. After receiving all of the relevant information, participants provided written informed consent before participating in the experiment.

### Participants

We recruited 16 gymnasts (11 men, 5 women, aged 18–24 years) and 16 healthy adults (8 men, 8 women, aged 19–22 years) as a non-gymnast group. All participants were right-handed. All gymnasts had been practicing at least for 7 years (range: 7–16 years) and had participated in an all-Japan intercollegiate gymnastic championship at least once. Note that two gymnasts were members of the Japanese national gymnastics team. The non-gymnasts group had no experience of gymnastic training. None of the participants had experienced any neurological and musculoskeletal disorders.

### Psychological assessments

#### Procedures

Motor imagery ability was tested using a psychological questionnaire translated into Japanese: The Kinesthetic and Visual Imagery Questionnaire (KVIQ-20) (21,26). Briefly, the KVIQ-20 tests how vividly a person is able to imagine their own movements subjectively, using two types of motor imagery: KMI and visual motor imagery (VMI). KMI involves imagining the feeling when we perform actual motor tasks. Conversely, VMI involves imagining to see ourselves or the field of vision where we perform the tasks (21,27). Participants sat comfortably in a chair next to the examiner and watched the examiner’s example once. Then, they actually performed the exercise, followed by VMI or KMI of the exercise they had just performed. Participants were then asked to evaluate the vividness of the motor imagery on a five-point ordinal scale (the more vivid the motor imagery, the higher the scale score). This procedure was repeated for 10 different simple exercises: neck flexion/extension, shoulder elevation, forward shoulder flexion, elbow flexion/extension, thumb-fingers opposition, forward trunk flexion, knee extension, hip abduction, foot tapping, and foot external rotation.

#### Analyses

We evaluated the vividness of motor imagery by summing all KVIQ scores for KMI and VMI, respectively. If a participant could imagine their movements perfectly, the score was 50 points.

### Physiological assessments

#### Recordings

Scalp EEG signals were recorded with eight passive Ag/AgCl electrodes around the sensorimotor area related to right upper limbs (Cz, C1, C3, C5, FC1, FC3, CP1, and CP3) in accord with the extended international 10-20 system. Electrodes with a diameter of 18 mm were mounted on an electrode cap (g.GAMMAcap 1027; Guger Technologies, Graz, Austria). Reference and ground electrodes were placed on the right and left earlobes, respectively. Surface electromyogram (EMG) signals were recorded from the right deltoid muscle (DEL) and the right extensor carpi radials muscle (ECR). Two passive Ag/AgCl electrodes with a diameter of 10 mm were placed over each muscle belly with inter-electrode distances of 20 mm. All EEG and EMG signals were amplified and bandpass-filtered (EEG, 0.5–1000 Hz; EMG, 2–1000 Hz) using a linked biosignal recording system (g.BSamp 0201a; Guger Technologies, Graz, Austria). All analog EEG and EMG signals were converted to digital signals at a sample rate of 1000 Hz using an AD converter with 16-bit resolution (NI USB-6259, National Instruments, Austin, TX, United States) that was controlled by data-logger software originally designed using MATLAB software (The MathWorks, Inc., Antic, MA, United States).

#### Procedures

Following the psychological assessment, we performed physiological EEG and EMG measurements. The participants sat comfortably in the seat. A computer monitor for visual feedback was placed 2 m in front of participants’ eyes. First, resting-state EEG was recorded over 60 s. Participants relaxed and fixated their eyes on a cross (+) displayed at the center of the monitor during recording. Participants then performed several practice trials of maximal voluntary contractions (MVCs) of wrist dorsiflexion and shoulder abduction. After these movements were practiced, participants performed MVC once each for wrist dorsiflexion and shoulder abduction. When performing each MVC, EMG activity of the contracting muscle was recorded. The EMG signals were full-wave rectified. We found a 0.5-s period of stable force exertion during MVC and calculated the integrated EMG value (iEMGmax) in this period. In the following experiment, 20% of this iEMGmax value was used as a target value for visual feedback.

The physiological data recordings during motor execution and KMI were performed after several practice trials. Visual feedback was presented on the screen, with a red cursor to represent muscle contraction level as a relative value in %iEMGmax and a vertical blue line to represent a target value. Furthermore, instructions for each phase, including “Rest”, “Relax”, “Ready”, “Contraction”, or “Imagery”, were displayed on the monitor. In the wrist dorsiflexion task, participants’ dominant hand was positioned on the armrest and fixed by a belt with the palm down. In the shoulder abduction task, the dominant upper limb was lowered to the side of the body while bending the elbow lightly, and the arm was fixed by a belt. In both the wrist dorsiflexion and shoulder abduction tasks, participants performed repeated motor execution and KMI according to the procedure used in our previous study (28).

The experimental paradigm is shown in Figure 1. In detail, each trial was started from the rest phase and the word “Rest” was displayed on the monitor for 7 s. During the rest phase, participants were able to adjust their posture freely and/or blink their eyes strongly. After the rest phase, the word “Relax” was displayed on the monitor for 3 s. During the relax phase, participants were instructed to relax as much as possible, without performing any movement. The word “Ready” was then displayed for 3 s, accompanied by a short sound presented every second. During the ready phase, participants prepared for the next instruction. After the ready phase, the word “Contraction” was displayed for 5 s. During the contraction phase, participants performed isometric voluntary contraction (wrist dorsiflexion or shoulder abduction) at 20% of iEMGmax by their dominant hand. In the contraction phase, participants were instructed to contract their muscle so that the cursor could follow the target line as accurately as possible. After the contraction phase, the word “Relax” was displayed for 2 s. In this relax phase, participants were instructed to relax as much as possible, without any movement. After the rest, relax, and ready phases, the imagery phase was started, and participants performed KMI of the preceding contraction for 5 s with their eyes open, and without any movement. During the imagery phase, we checked that no EMG activity occurred. When the imagery phase finished, the relax phase was presented again for 2 s. This flow was conducted for each trial, including motor execution and KMI, and five trials were repeated within each set. Six sets were performed for each task. Thus, a total 30 trials were performed for both wrist flexion and shoulder abduction tasks. We set the wrist dorsiflexion and the shoulder abduction task in a randomized order across participants. The duration of the set interval was longer than 2 min, to provide sufficient rest for participants.

**Figure 1.**
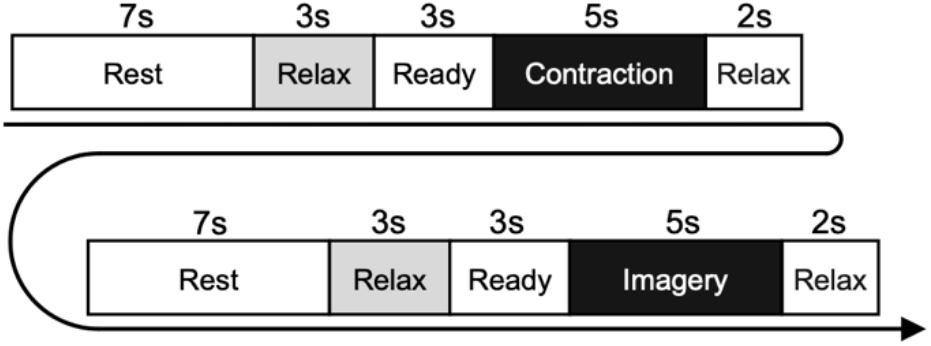
Experimental paradigm of physiological experiment. Participants performed isometric contraction in the contraction phase and performed motor imagery of the same movement in the imagery phase. The diagram shows the flow in each trial, which was repeated five times within each set. Six sets were performed for each of the wrist dorsiflexion and shoulder abduction tasks.

#### Analyses

To remove noise arising from the electric power, the EEG and EMG signals were notch-filtered at 50 Hz. The EEG signals over C1 and C3 were derived with a four-neighbor Laplacian spatial filter. For example, in the case of C3, the EEG signal over C3 was subtracted by an average of C1, C5, FC3 and CP3. The Laplacian derivation method is known to strongly emphasize cortical activity originating below the electrode of interest (29). If Laplacian-derived EEG included potentials exceeded 50 μV, we considered the trial to contain an artifact, and excluded the data from future analyses. Additionally, visual inspection was performed to reject additional artifacts missed by the automatic inspection.

ERDs during motor execution and KMI of each task were calculated as follows. After separating the data into motor execution and KMI periods, we extracted the 30 1-s data windows in the same period from the data for each trial. Then, fast Fourier transformation was performed using Welch’s method for the data (window length, 1 s; window function, Hanning-window; overlap, 0), and the power spectrum densities (PSDs) of the EEG signal were calculated. This process was repeated by sliding the 1 s data window in 50 ms steps. The ERDs were calculated using the following equation:

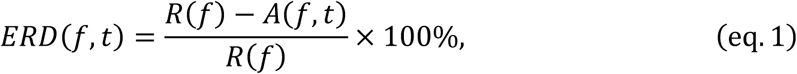

where *A* is the EEG PSDs at time *t*, frequency *f*, and *R* is the mean PSDs of the baseline period (last 1 s in the relax phase). This equation indicates that the positively greater the ERD value, the larger the decrease in EEG PSD during motor execution or KMI compared with the relax phase. Because the most reactive frequency band of ERD was slightly different across participants (30), we determined an electrode and the 3 Hz-frequency width showing the largest ERD in each of the α-band (8–15 Hz) and β-band (16–35 Hz) during motor execution. Because it has been suggested that the functional roles played by ERD differ between the α- and β-bands (31,32), we analyzed ERDs from these two frequency bands separately. The magnitude of ERD in the α-band (αERDmax) and β-band (βERDmax) were measured by calculating the peak value of ERD for motor execution and KMI of each task, respectively (28,33).

To compare the features of EEG during the relax phase in the task with continuous resting-state for a prolonged period, we also analyzed the α-band or β-band PSDs for both data sets. For the relax phase EEG, the final 1 s periods in the relax phase, which were used as the baseline periods for ERD analyses, were extracted from all trials, and combined to create a 60 s relax phase EEG signal. For the resting-state EEG, a continuous 60 s period with few artefacts was extracted. We then calculated the ratio of the sum of EEG power within the α-band or β-band PSD to that of the entire frequency range (4–50 Hz) (named, EEGα-PSD and EEGβ-PSD) for both data sets, and compared these values between relax phase EEG during tasks and resting-state EEG.

### Statistical analyses

Two-side unpaired t-tests were performed on VMI and KMI scores of KVIQ between groups (non-gymnasts vs. gymnasts), to confirm differences in the subjective vividness of motor imagery between them. To test differences in αERDmax and βERDmax during motor execution or KMI, we performed two-way mixed-model analysis of variance (ANOVA) between participant groups (gymnasts and non-gymnasts) and tasks (wrist dorsiflexion and shoulder abduction). If the interaction was significant, we performed a two-sided unpaired t-test for groups (gymnasts vs. non-gymnasts) and a two-sided paired t-test for tasks (wrist dorsiflexion vs. shoulder abduction). To test differences in the EEGα-PSD or EEGα-PSD between resting-state EEG and relax phase EEG during tasks, we also performed two-way ANOVA between participant groups (gymnasts and non-gymnasts) and conditions (resting-sate and relax phase). The p-values of 0.05 were used to indicate statistical significance. All statistical analyses were performed using SPSS statistics software (IBM SPSS Statistics 25, IBM developerWorks, Tokyo, Japan).

## Results

### KVIQ

Figure 2 shows group data (mean ± S.D.) for KVIQ scores obtained from VMI and KMI tasks between gymnasts and non-gymnasts. The KVIQ scores were significantly greater in gymnasts, both in VMI (gymnasts, 42.68 ± 6.22; non-gymnasts, 35.44 ± 8.73, *p* = 0.011) (Figure 2A) and KMI (gymnasts, 43.50 ± 6.78; non-gymnasts, 35.94 ± 8.24, *p* = 0.008) (Figure 2B). The KVIQ results indicate that gymnasts subjectively evaluated how vividly they could imagine their own movements.

**Figure 2.**
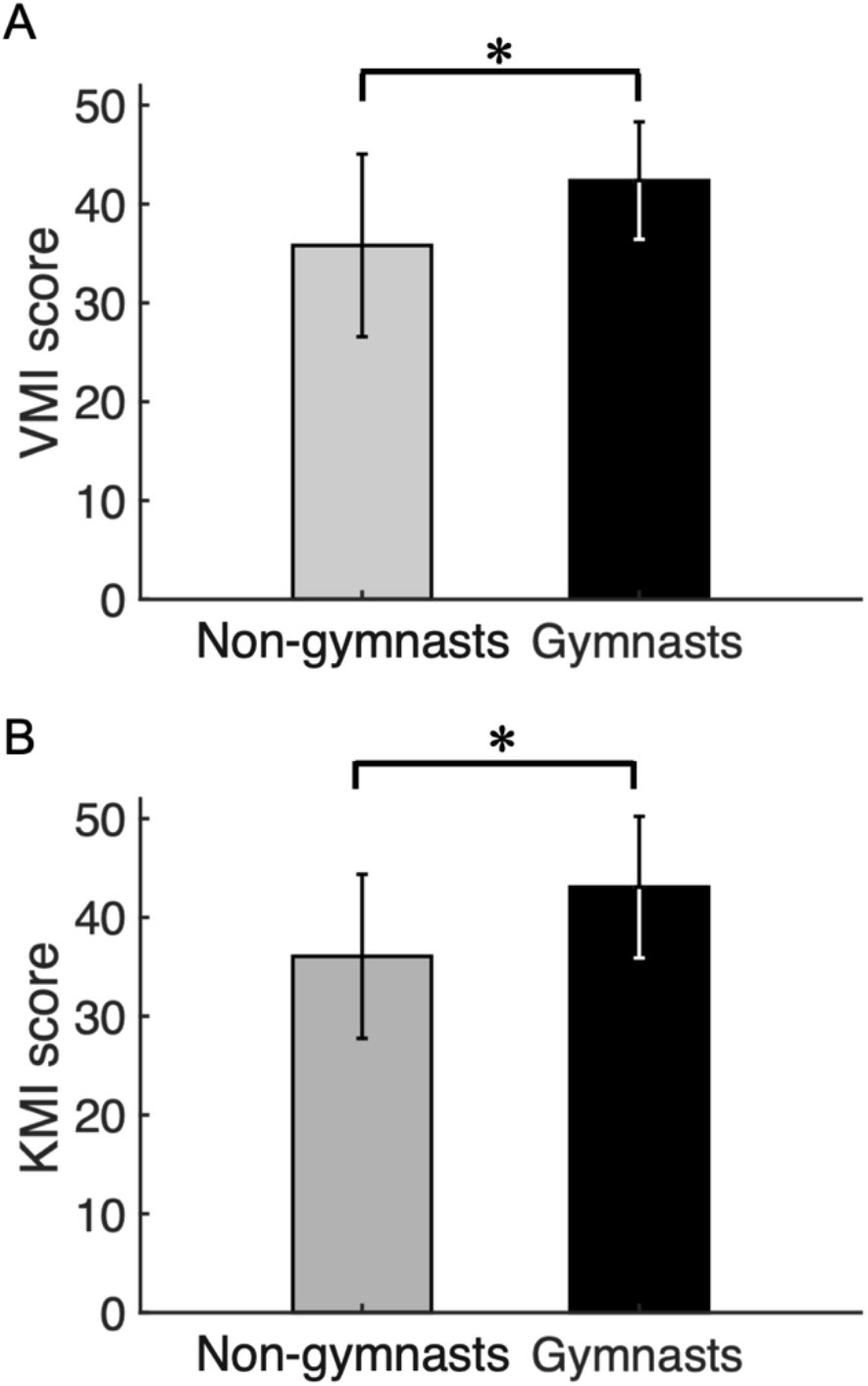
Results of psychological experiment. Group data (Mean ± S.D.) for visual motor imagery (VMI) (A) and kinesthetic motor imagery (KMI) scores (B) obtained from the Kinesthetic and Visual Imagery Questionnaire (KVIQ) are shown for both groups. The gray bars represent data for non-gymnasts, while the black bar represents data for gymnasts. *P < 0.05.

### ERD magnitude

Typical examples of EEG signals, EEG time-frequency maps and ERD time courses during wrist dorsiflexion from a non-gymnast and a gymnast are shown in Figure 3A and 3B, respectively. From these time-frequency maps, a decrease of EEG power can be observed around 12 Hz and 22 Hz in the contraction phase (0 to 5 s) compared with the relax phase (−6 to −3 s) in both participants when performing motor execution.

**Figure 3.**
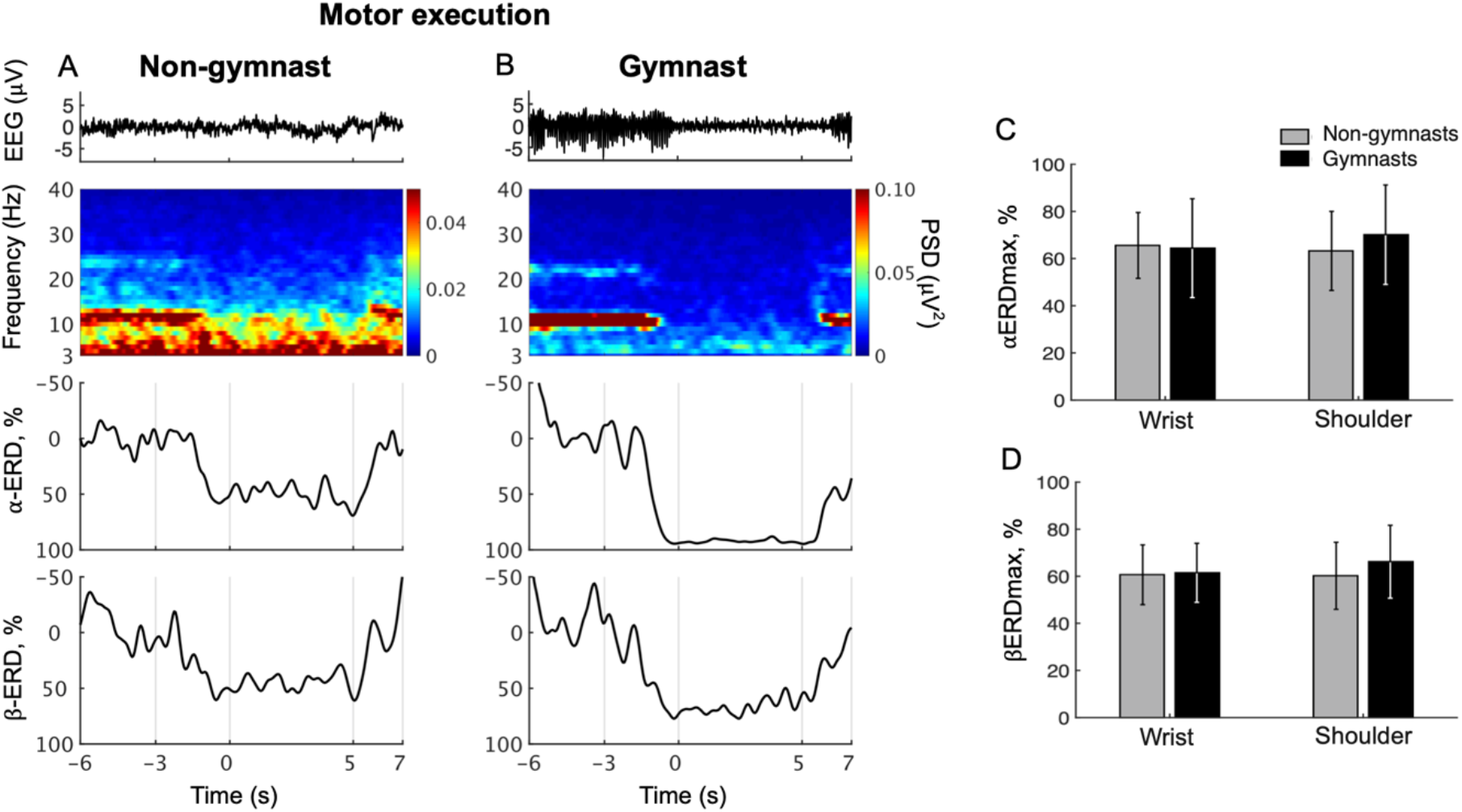
Results from physiological experiments for motor execution. Typical time courses of single trial electroencephalogram (EEG), time-frequency map, α-band event-related desynchronization (ERD), and β-band in wrist dorsiflexion motor execution are shown for non-gymnast (A) and gymnast (B) participants. Note that participants performed wrist dorsiflexion motor execution from 0 s to 5 s. Group data (Mean ± S.D.) for the maximal peak of ERD both in the α- (αERDmax) (C) and β-bands (βERDmax) (D) during motor execution are shown across groups and tasks. The gray bars represent data for non-gymnasts, while the black bar represents data for gymnasts. No significant differences were observed across groups and tasks.

Figure 3C shows group data for αERDmax during wrist dorsiflexion and shoulder abduction motor execution. An ANOVA on αERDmax during motor execution revealed no significant effects of group (F_1,30_ = 0.209, p = 0.651) and task (F_1,30_ = 0.831, p = 0.369), while a significant interaction was obtained (F_1,30_ = 4.654, p = 0.0391). An unpaired t-test for group revealed no significant difference in αERDmax between gymnasts and non-gymnasts for wrist dorsiflexion (gymnasts, 64.41 ± 20.96; non-gymnasts, 65.58 ± 13.97, p = 0.855) and shoulder abduction task (gymnasts, 70.13 ± 21.07; non-gymnasts, 63.26 ± 16.77, p = 0.316). A paired t-test for task revealed no significant difference in αERDmax between wrist dorsiflexion and shoulder abduction execution both for non-gymnasts (wrist dorsiflexion, 65.58 ± 13.97; shoulder abduction, 63.26 ± 16.77, p = 0.303) and gymnasts (wrist dorsiflexion, 64.41 ± 20.96; shoulder abduction, 70.13 ± 21.07, p = 0.078). Figure 3D shows group data for βERDmax during wrist dorsiflexion and shoulder abduction motor execution. An ANOVA on the βERDmax during motor execution with groups and task revealed no significant effects of group (F_1,30_ = 0.571, p = 0.456) and task (F_1,30_ = 1.249, p = 0.273), and interaction (F_1,30_ = 1.815, p = 0.188). The results revealed no effects of sports experience and body part on ERD magnitude during motor execution.

Typical examples of EEG signals, EEG time-frequency maps and ERD time courses during wrist dorsiflexion KMI from a non-gymnast and a gymnast are shown in Figure 4A and 4B, respectively. From the time-frequency maps for a non-gymnast participant, we did not observe clear ERD in the imagery phase (0 to 5 s) compared with the relax phase (−6 to −3 s) in both the α-band and β-band. (Figure 4A). Conversely, clear ERD can be observed in the time-frequency-map for the gymnast participant around 12 Hz and 22 Hz (Figure 4B).

**Figure 4.**
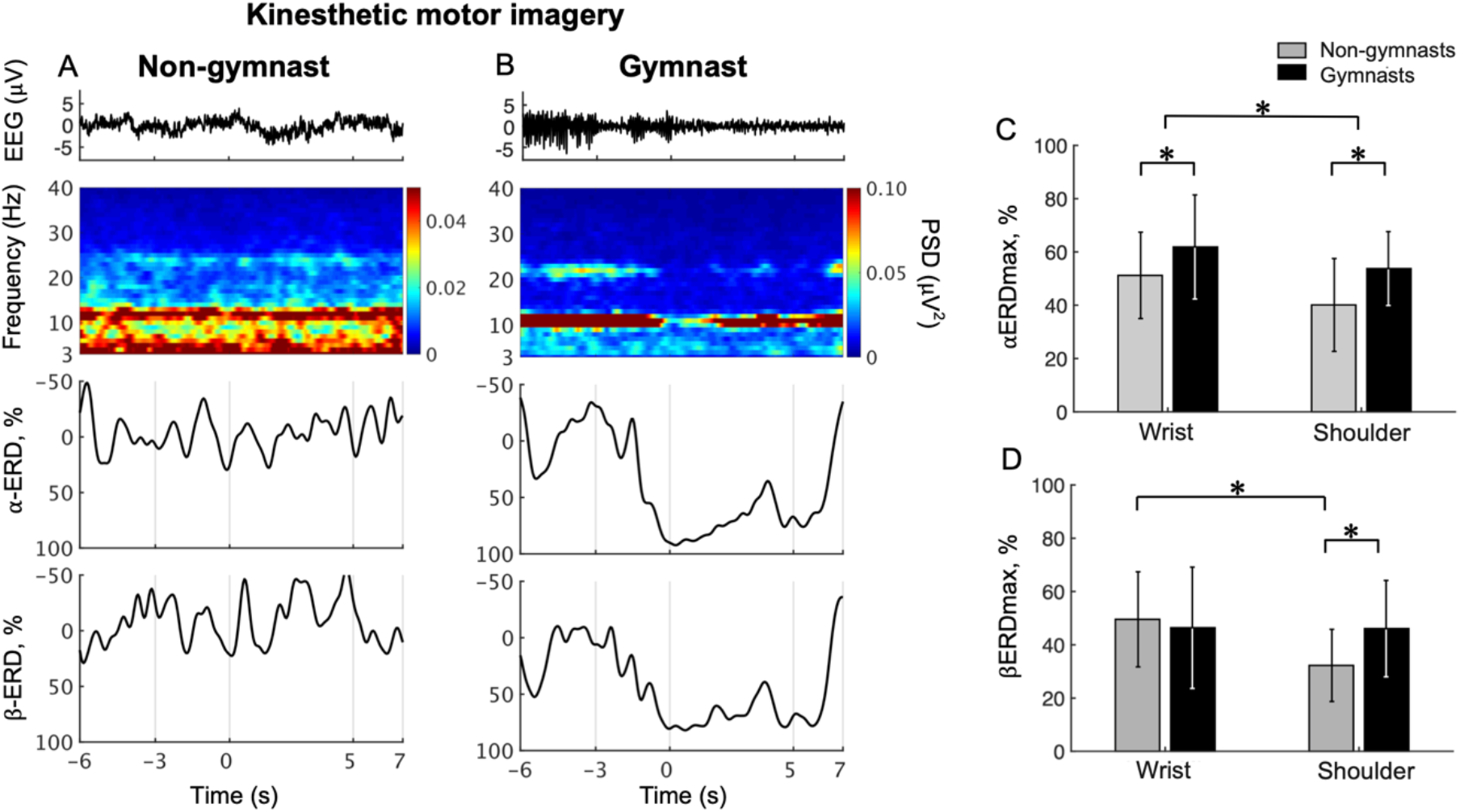
Results from physiological experiments for KMI. Typical time courses of single trial EEG, time-frequency map, α-band ERD, and β-band in wrist dorsiflexion KMI are shown for non-gymnast (A) and gymnast (B) participants. Note that participants performed wrist dorsiflexion KMI from 0 s to 5 s. Group data (Mean ± S.D.) for αERDmax (C) and βERDmax (D) during KMI are shown across groups and tasks. The gray bars represent data for non-gymnasts, while the black bar represents data for gymnasts. *P < 0.05.

Figure 4C shows group data for the αERDmax during wrist dorsiflexion and shoulder abduction KMI. An ANOVA on αERDmax during KMI showed significant effects of group (F_1,30_ = 5.437, p = 0.027) and task (F_1,30_ = 10.975, p = 0.002). No significant interaction effect (F_1,30_ = 0.266, p = 0.610) was observed. Figure 4C shows group data for βERDmax during wrist dorsiflexion and shoulder abduction KMI. An ANOVA on βERDmax during KMI showed no significant effects of group (F_1,30_ = 0.876, p = 0.357), but significant effects of task (F_1,30_ = 8.019, p = 0.008) and a significant interaction (F_1,30_ = 7.421, p = 0.010). An unpaired t-test for group revealed a significant difference in βERDmax between gymnasts and non-gymnasts for the shoulder abduction task (gymnasts, 46.08 ± 18.09; non-gymnasts, 32.27 ± 17.84, p = 0.021) but not for the wrist dorsiflexion task (gymnasts, 46.42 ± 22.76; non-gymnasts, 49.56 ± 17.84, p = 0.666). A paired t-test for task revealed a significant difference in βERDmax between the wrist dorsiflexion and shoulder abduction KMI conditions for non-gymnasts (wrist dorsiflexion, 49.56 ± 17.84; shoulder abduction, 32.27 ± 17.84, p = 0.002) but not for gymnasts (wrist dorsiflexion, 46.42 ± 22.76; shoulder abduction, 46.08 ± 18.09, p = 0.938). These results indicated that gymnastics experience affected ERD magnitude during KMI of sports non-specific movements.

### Comparison of EEGα-PSD and EEGβ-PSD between resting-state EEG and relax phase EEG during tasks

Figure 5A shows group data for the EEGα-PSD in resting-state EEG and relax phase EEG during the tasks. An ANOVA examining EEGα-PSD data revealed no significant effects of group (F_1,30_ = 0.486, p = 0.491) or condition (F_1,30_ = 4.135, p = 0.051); however, a significant interaction (F_1,30_ = 6.382, p = 0.017) was observed. An unpaired t-test for group revealed no significant difference in EEGα-PSD during resting-state EEG (gymnasts, 0.473 ± 0.155; non-gymnasts, 0.482 ± 0.166, p = 0.878) and that during the relax phase (gymnasts, 0.482 ± 0.155; non-gymnasts, 0.403 ± 0.125, p = 0.878; p = 0.122) between gymnasts and non-gymnasts. In non-gymnasts, a paired t-test revealed significant differences in the EEGα-PSD between conditions (resting-state, 0.482 ± 0.166; relax phase, 0.403 ± 0.125, p = 0.008). However, in gymnasts, no significant differences in EEGα-PSD were observed between conditions (resting-state, 0.473 ± 0.155; relax phase, 0.481 ± 0.155, p = 0.715). Figure 5B shows group data for EEGβ-PSD. An ANOVA on EEGβ-PSD showed no significant effects of group (F_1,30_ = 0.106, p = 0.747) and condition (F_1,30_ = 1.787, p = 0.191), and no significant interaction (F_1,30_ = 0.063, p = 0.804). These results indicate that, in non-gymnasts, the EEGα-PSD was smaller in the relax phase EEG than during resting-state EEG, while such a difference was not observed in gymnasts. Conversely, EEGβ-PSD did not differ between resting-state EEG and relax phase EEG in both gymnasts and non-gymnasts.

**Figure 5.**
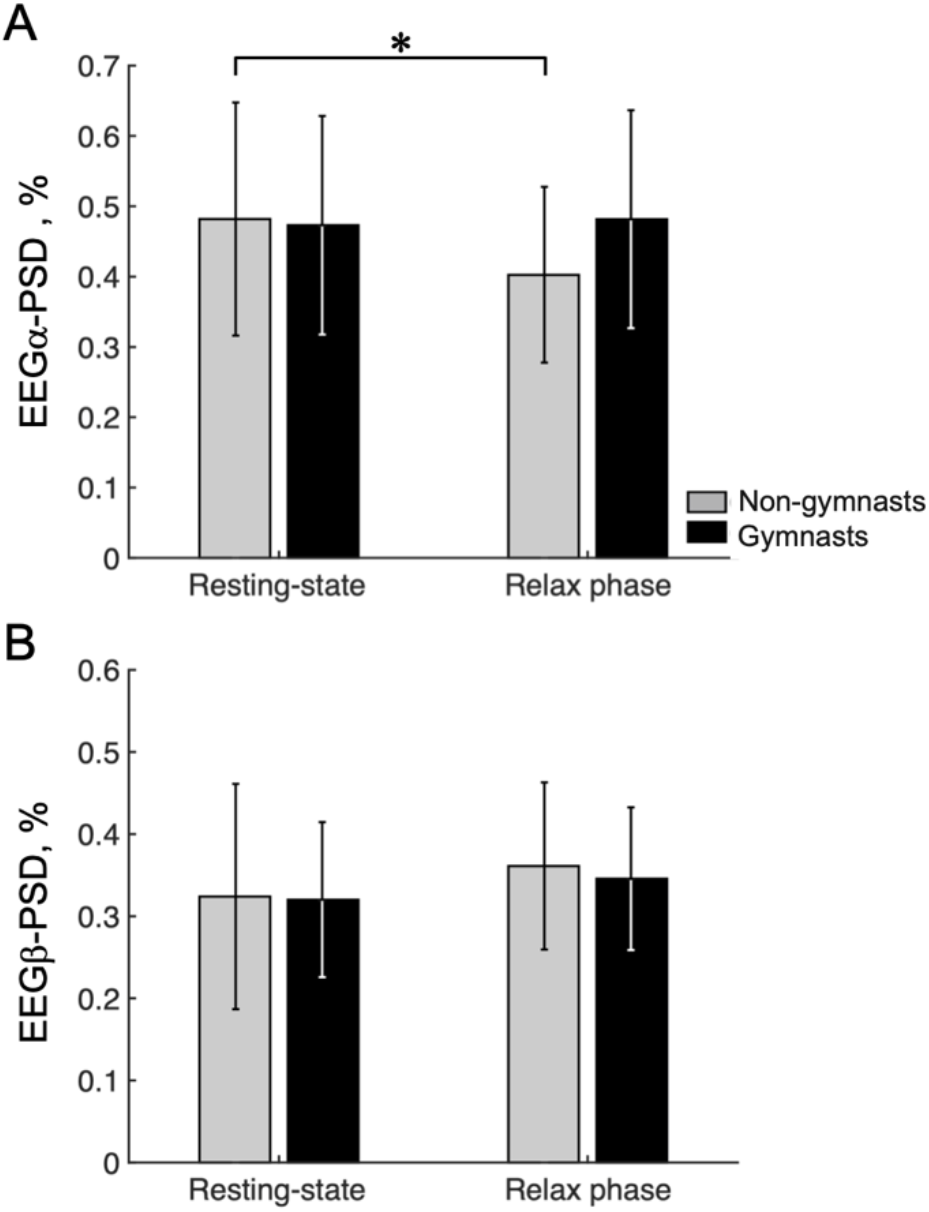
Results of the ratio of the α-band and β-band power spectrum densities. Group data (Mean ± S.D.) for the ratio of the α-band (A) and β-band power spectrum densities (PSD) (B) are shown for both groups. The gray bars represent data for non-gymnasts, while the black bar represents data for gymnasts. *P < 0.05.

## Discussion

The purpose of the present study was to clarify differences in bioelectric sensorimotor rhythm during KMI of sports non-specific movements between gymnasts and non-gymnasts. The results revealed that, when required to repeatedly switch between relaxing and motor execution or KMI of sports non-specific movements, the ERD magnitude during KMI was significantly greater in gymnasts, who subjectively evaluated their imagery including KMI as more vivid, while no difference between groups was observed during motor execution. In particular, the ERD magnitude in the α-band was greater in gymnasts compared with non-gymnasts, both in wrist dorsiflexion and shoulder abduction KMIs, whereas the ERD magnitude in the β-band was greater in gymnasts only in shoulder abduction KMI.

### Difference in KMI ability of sports non-specific movements between gymnasts and non-gymnasts

We evaluated ERD as a physiological indicator of KMI ability in the present study, because it is considered to reflect changes in corticospinal excitability (24) and is associated with subjective vividness of KMI measured by KVIQ (28). It should be noted that the present results revealed greater ERD magnitude during KMI of sports non-specific movements in gymnasts than in non-gymnasts, although differences in neural activity between athletes and non-athletes have not been reported in motor imagery of sports non-specific movements in previous studies using EEG (17), MEG (18), TMS (20) or fMRI (19). This may be related to the fact that gymnasts perform motor imagery including KMI frequently as a part of their daily practice to reduce the risk of serious injury in their practice. Furthermore, a previous psychological study showed that the vividness of motor imagery of sports non-specific movements was higher in athletes engaged in individual and/or non-contact sports compared with athletes engaged in team and/or contact sports (25). Thus, as gymnasts have superior motor imagery ability among athletes, they may provide a particularly suitable population for highlighting differences in neural activity during KMI of sports non-specific movements compared with non-athletes.

It is possible that the present task protocol, in which participants performed KMI following motor execution repeatedly in the order of seconds, led to the current finding of greater ERD magnitude in gymnasts. In psychological questionnaires, the conventional procedure for measuring motor imagery ability is to examine participants while they perform motor execution then motor imagery in one trial, and subjectively evaluate the vividness of the motor imagery of the preceding movement (21). In physiological experiments, however, the conventional procedure involves evaluating neural activity while participants perform only motor imagery (17–20). Thus, there has been a methodological gap in the approach for examining of motor imagery between psychological questionnaire studies and experimental physiological studies. In response to direct questioning in the current study, gymnasts reported that they usually perform actual movements and KMIs alternately in their daily practice. Therefore, the method for measuring KMI in the present physiological experiment was designed in accord with the procedure of psychological questionnaire measurement. The current physiological findings may have been due to differences between gymnasts and non-gymnasts in the ability to flexibly modulate corticospinal excitability when imagining their own movements, by referring to the actual movement.

As shown in equation (1), we were able to confirm that the ERD was determined by both the degree of synchronization during the relax phase (*R(f)*) and the degree of desynchronization during KMI (*A(f, t)*). As shown in Fig. 5, first, gymnasts could return their sensorimotor α-rhythm during the relax phase in the task to the same power level as during the resting-state for 60 s, whereas non-gymnasts could not. Thus, gymnasts appeared to be good at relaxing deeply by making their sensorimotor rhythms more synchronized within a short period of time. However, higher EEGα-PSD during the relax phase does not appear to be the only factor involved in gymnasts’ greater ERD magnitude in the α-band. As shown in Fig. 2, second, differences in αERDmax between groups were not observed in motor execution but were found in KMI. Thus, gymnasts also appeared to be good at increasing corticospinal excitability by making their sensorimotor rhythms more desynchronized, even in KMI. Overall, the present results demonstrate that gymnasts have the ability to generate a clear contrast in the state of the sensorimotor cortex, when required to repeatedly switch across relaxing, motor execution and KMI conditions. On the basis of the current findings, we believe that the ability to modulate the brain state without any movement is a core aspect of superior KMI ability in gymnasts.

### Difference in functional role of ERD between α- and β-band

Interestingly, the present study demonstrated different results between ERD magnitude in the α-band and β-band. Several previous studies reported that functional roles played by the sensorimotor rhythm are different between frequency bands. During actual muscle contraction with weak-to-moderate intensity, the sensorimotor rhythm is known to be coherent with EMG activity only in the β-band, with no significant coherence in the α-band (34–36). When focusing on oscillatory power itself, the EEG spectral power in the sensorimotor area contralateral to the contracted/imagined limb was decreased in both the α- and β-bands, while that in task-irrelevant cortical regions was increased in the α-band (22,37), but not in the β-band (31). During KMI, the ERD magnitude was increased by increasing task demand in the β-band, but not in the α-band (32,38). Taken together, these findings suggest that the functional roles of sensorimotor rhythms for movement/imagery should be distinguished between the α-band and β-band.

First, αERDmax was larger in gymnasts than non-gymnasts during both KMI tasks (i.e., wrist dorsiflexion and shoulder abduction). In task-relevant cortical regions, neural populations are assumed to be disinhibited by the ERD of the sensorimotor area in the α-band, which would allow reallocation of computational resources (32). However, task-irrelevant cortical regions are assumed to be inhibited by enhancing their α-oscillations (23). As gymnasts are required to perform skilled movements successively, they are trained to quickly switch their attention across their body parts by facilitating task-relevant regions and inhibiting task-irrelevant regions. The present results regarding ERD in the α-band would reflect such an ability of gymnasts.

Second, βERDmax was larger in gymnasts only during shoulder abduction KMI, but not during wrist dorsiflexion KMI. This task-specificity in βERDmax may be caused by ERD in the β-band playing a role in the calculation of specific motor commands. In general, wrist movement is used frequently in daily life, which makes it easy for most people to perform wrist dorsiflexion KMI. Thus, βERDmax would not be differed between gymnasts and non-gymnasts in the wrist dorsiflexion KMI task. However, as isometric shoulder abduction is a movement rarely used in daily life, it may be difficult for most people to perform this KMI. Conversely, gymnasts are well-trained to move their upper limbs, including the shoulder joints, both dynamically (i.e., giant swing) and statically (i.e., handstand and rings). Therefore, it would be easy for gymnasts to imagine shoulder abduction because they are skilled at adjusting the movement parameters of their shoulder joints. We assume that ERD in the β-band is an indicator for how precisely a person can imagine their own movement kinesthetically.

### Limitations

In the present study, only upper limb movements (i.e., wrist dorsiflexion and shoulder abduction) were examined. Gymnasts use their upper limb muscles specifically as anti-gravity muscles for postural control, such as handstand and pommel horse. This usage of the upper limbs is unique relative to the movements of non-gymnasts. The uniqueness of gymnasts’ upper limb usage may lead to superior KMI ability regarding upper limb movements. Thus, we cannot clearly predict whether similar results would be obtained when performing similar experiments for other body parts. However, the KVIQ results demonstrated that gymnasts tended to show higher scores for all movements. In addition, most gymnasts perform motor imagery of various body parts in their daily practice. On the basis of these findings, we speculate that gymnasts have superior KMI ability irrespective of body parts, although confirming this possibility will require further investigation.

The present study is the first to observe differences in physiological indices between athletes and non-athletes. This means that the present study can bridge the gap between psychology and physiology studies regarding differences in KMI ability of sports non-specific movements between athletes and non-athletes. However, because only gymnasts participated in this study, it is unclear whether the present results are specific to gymnasts or apply generally to athletes performing any sports. Because differences in motor imagery ability of sports non-specific movements would be expected among athletes, further investigation is needed to elucidate sports-specific differences in motor imagery ability for sports non-specific movements. In any case, the present study indicated the importance of comparing corticospinal excitability measured by ERD for evaluating KMI ability. In future studies, imagery training using bioelectrical signals may provide a useful tool for improving the motor imagery ability of athletes.

## Conclusion

The present study demonstrated that, during KMI of sports non-specific movements, the corticospinal excitability measured by ERD magnitude was significantly greater in gymnasts compared with non-gymnasts. These results are consistent with higher subjective vividness of KMI in gymnasts measured using the KVIQ psychological questionnaire. The observed signature of flexibly modulating sensorimotor rhythm with no movement would be the basis of their superior KMI ability of sports non-specific movements in gymnasts.

## Author Contributions

H.S. and J.U. conceptualized and designed the study, interpreted data, wrote the manuscript, and acquired fundings. H.S. acquired and analyzed data. J.U. supervised the study.

## Acknowledgements

This work was supported by grants from the Grant-in-Aid for Scientific Research (B) (Japan Society for the Promotion of Science, JSPS) (grant number 20H04091) to JU, a designated donation from Living Platform, Ltd, Japan to JU, and Taikichiro Mori Memorial Research Grants to HS. We thank Ms. Tomomi Hamaoka, Ms. Kana Iijima, and Ms. Chieko Matsuda for their secretarial assistance, and Mr. Hisato Toriyama, Mr. Ryoichiro Yamazaki, Mr. Takuya Ideriha, Ms. Rina Suzuki and all other members of our laboratory for their useful comments on the work. We thank Mr. Hisashi Mizutori for his practical comments on the work. We thank Benjamin Knight, MSc., from Edanz (https://www.jp.edanz.com/ac), for editing a draft of this manuscript.

For all authors, no conflicts of interest were declared.

